# Transcriptional regulatory network controlling the ontogeny of hematopoietic stem cells

**DOI:** 10.1101/856559

**Authors:** Peng Gao, Changya Chen, Elizabeth D. Howell, Yan Li, Joanna Tober, Yasin Uzun, Bing He, Long Gao, Qin Zhu, Arndt Siekmann, Nancy A. Speck, Kai Tan

## Abstract

Hematopoietic stem cell (HSC) ontogeny is accompanied by dynamic changes in gene regulatory networks. We performed RNA-Seq and histone mark ChIP-Seq to define the transcriptomes and epigenomes of cells representing key developmental stages of HSC ontogeny in the mouse. The five populations analyzed were embryonic day 10.5 (E10.5) endothelium and hemogenic endothelium from the major arteries (dorsal aorta, umbilical and vitelline), an enriched population of pre-hematopoietic stem cells (pre-HSCs), fetal liver HSCs, and adult bone marrow HSCs. We observed dynamic and combinatorial epigenetic changes that mark regulatory DNA sequences including gene promoters and enhancers. Using epigenetic signatures, we identified enhancers for each developmental stage. Only 12% of enhancers are primed, and 78% are active, suggesting the vast majority of enhancers are established *de novo* at the developmental stages where they are required to control their target genes, without prior priming in earlier stages. We constructed developmental-stage-specific transcriptional regulatory networks during HSC ontogeny by linking enhancers and predicted bound transcription factors to their target promoters using a novel computational algorithm. Our computational analyses predicted known transcriptional regulators for the endothelial-to-hematopoietic transition, validating our overall approach, and identified putative novel transcription factors whose regulon activities correlate with the emergence of pre-HSCs. We validated roles for the broadly expressed transcription factors SP3 and MAZ in arterial hemogenic endothelium. Our data and computational analyses provide a useful resource for uncovering regulators of HSC formation.

## Introduction

Hematopoietic stem and progenitor cells (HSPCs) differentiate from a small population of endothelial cells in the embryo called hemogenic endothelium (HE) (Chen et al., 2009; Zovein et al., 2008). The process of HSPC formation from HE involves an endothelial to hematopoietic transition (EHT), in which HE, consisting of flat cells in a monolayer interconnected by tight junctions, undergoes a transition to form round cells that detach from the endothelial layer, and enter circulation (Bos et al., 2015; Kissa and Herbomel, 2010). HE is found at multiple anatomic sites in the embryo such as the yolk sac, the major caudal arteries including the vitelline and umbilical arteries, and the dorsal aorta where it is flanked by the developing urogenital ridges in the so-called aorta-gonad-mesonephros (AGM) region (North et al., 1999). The first HE cells in the mouse appear in the yolk sac at approximately embryonic day (E) 8.5, they are most abundant in the major arteries at E9.5, and the majority of HE cells in the major arteries undergo EHT between E9.5 and E12.5(North et al., 1999; Tober et al., 2013; Yokomizo and Dzierzak, 2010). The differentiation of HSPCs from HE in the major arteries in most vertebrate organisms involves an intermediate step in which the HSPCs briefly accumulate in clusters attached to the luminal side of the arterial wall, and are then released into the circulation to colonize the fetal liver (FL).

Although the intra-arterial hematopoietic clusters (IACs) in mouse and human embryos contain hundreds of HSPCs, only 1-3 of them are functional adult-repopulating HSCs (Ivanovs et al., 2011; Kumaravelu et al., 2002; Yokomizo and Dzierzak, 2010). IACs in mouse embryos also contain between ∼150 (as measured by *ex vivo* maturation followed by transplantation) to ∼600 (determined by multicolor lineage tracing) immature precursors of HSCs (pre-HSCs) that mature into HSCs either within or prior to colonizing the FL (Ganuza et al., 2017; Kieusseian et al., 2012; Rybtsov et al., 2016; Taoudi et al., 2008). FL HSCs ultimately take up permanent residence in the bone marrow (BM) where they mature into adult BM HSCs (Bowie et al., 2007). In summary, the following five hematopoietic populations represent a continuum of embryonic to adult HSC maturation: endothelium (Endo) > HE > pre-HSCs > FL HSCs > BM HSCs.

Global epigenetic profiling has provided important insights into the molecules and mechanisms that regulate HSPC formation from endothelium. Most epigenetic studies of HSPC formation have been performed on cells generated *ex vivo* from mouse and human embryonic stem (ES) cells (Obier and Bonifer, 2016). For example, chromatin immunoprecipitation followed by sequencing (ChIP-seq) for histone marks and DNase followed by sequencing (DNase-seq), combined with ChIP-seq analysis of transcription factor (TF) occupancy were used to define the chromatin landscape and predict transcriptional regulatory networks at each step of the differentiation process from ES cells to adherent macrophages (Goode et al., 2016). This comprehensive study identified potential regulators of specific steps in this developmental trajectory, such as the TEAD/YAP factors, that were validated in functional assays. However, ES cell cultures are thought to model yolk sac HE and HSPC formation (Vanhee et al., 2015), whereas HSCs in the adult mouse are derived from the major arteries in the embryo from a distinct population of HE (Chen et al., 2011; Slukvin and Uenishi, 2019). Little is known about the dynamics of the epigenome during HSC formation from the major arteries since many of the early precursors are extremely rare, precluding the application of most ChIP-Seq protocols. There are good reasons to expect that HSPC formation in the yolk sac and major arteries will involve overlapping but distinct transcriptional regulatory networks. It is known, for example, that HSPC formation from the major arteries requires Notch signaling and the TF MYB, while HSPC formation in the yolk sac requires neither (Hadland et al., 2004; Mucenski et al., 1988; Tober et al., 2008). Our previous bulk RNA-seq comparison of arterial and yolk sac HE revealed a large number (∼2000) of differentially expressed genes (DEGs) (Gao et al., 2018). We predict that distinct requirements for HSPC formation from arterial endothelium are layered on top of a canonical EHT pathway that is required in all endothelial cells.

Here we analyze the transcriptomes, and with a low-cell-number ChIP-Seq protocol profile the genomic locations of four histone modification marks over the course of HSC ontogeny from arterial endothelium. We also develop an algorithm for predicting transcriptional regulatory networks operational at discrete stages of HSC formation, and demonstrate the activity of two TFs, SP3 and MAZ, during EHT.

## Results

### Purification and characterization of cell populations

We profiled five populations of cells spanning the continuum from endothelium to BM-HSCs (Figure 1). These include HE and Endo from the major arteries (dorsal aorta, umbilical and vitelline) of E10.5 embryos, separated based on their expression (or lack of expression) of green fluorescent protein (GFP) from the endogenous *Runx1* locus (Lorsbach et al., 2004), as described previously (Gao et al., 2018) (Figure S1). Both Endo and HE cells can form endothelial tubes in culture, whereas endothelial cells with the ability to form CD45^+^ blood cells in culture are highly enriched in the purified Runx1:GFP^+^ HE population (Gao et al., 2018). We also purified pre-HSCs, defined as cells that have newly differentiated from HE and are destined to become HSCs, but are not fully mature, and consequently cannot directly engraft adult recipients (Rybtsov et al., 2011) (Figure S1). All HSCs and pre-HSCs in the major arteries express a transgene from which GFP is expressed from the *Ly6a* (Sca1) regulatory sequences (de Bruijn et al., 2002; Ma et al., 2002; Tober et al., 2018). Only ∼15% of IAC cells are Ly6a:GFP^+^ (Li et al., 2014), thus by incorporating Ly6a:GFP into sorting strategies we enrich for pre-HSCs and HSCs within IAC cells. As pre-HSCs greatly outnumber HSCs at E11.5, for simplicity sake we will refer to the purified CD31^+^ CD144^+^ c-Kit^+^ Ly6a:GFP^+^ cells as pre-HSCs. Finally, we purified E14.5 FL HSCs (Lin^-^ Sca1^+^ c-Kit^+^ CD48^-^ CD150^+^) and adult BM HSCs (Lin^-^ Sca1^+^ c-Kit^+^ CD48^-^ CD150^+^ CD135^-^) (Figure S1). On average, we used 83,157 and 21,223 purified cells from each population for RNA-Seq and ChIP-Seq assays, respectively (Table S1 and S2).

**Figure 1.**
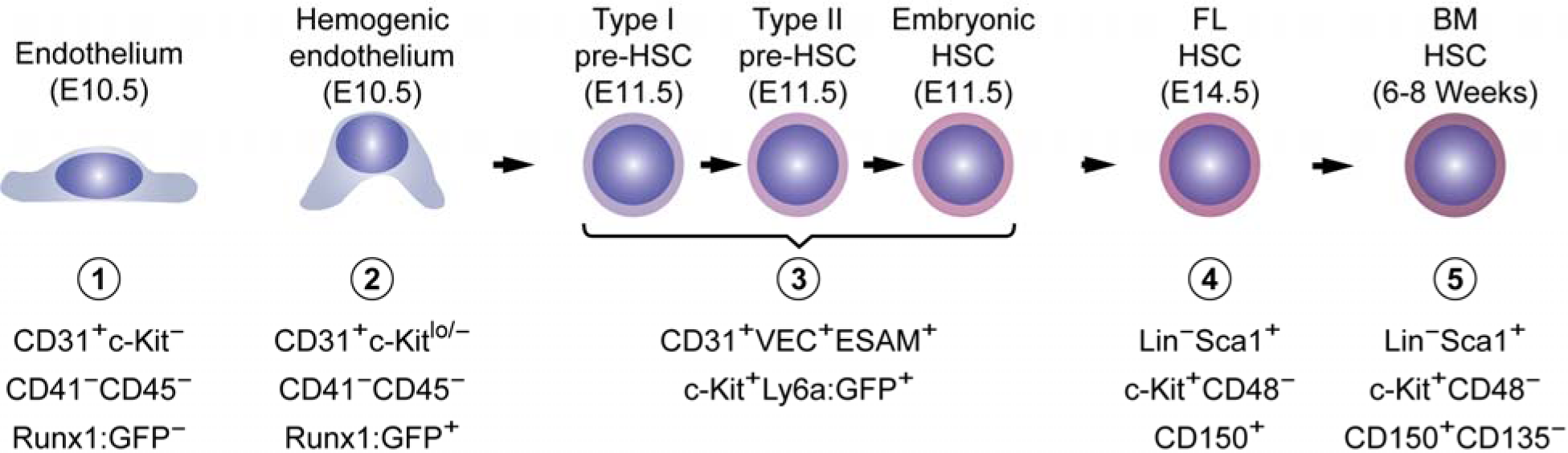
Purification of cells representing five stages of HSC ontogeny. The surface marker phenotypes of the cell populations purified are illustrated. Representative sort plots are presented in Figure S1, and functional characterization of the cells reported in Gao et al. (Gao et al., 2018).

### Transcriptome dynamics during HSC ontology

To identify changes in transcriptomes during HSC ontogeny, we performed RNA-Seq using sorted cells at the five developmental stages (Endo, HE, pre-HSC, FL HSC, BM HSC). The average correlation coefficient between biological replicates is 0.98 (Figure S2). We detected an average of 12,511 expressed genes at a FPKM threshold of 1, and 5,025 differentially expressed genes (DEGs) in each population with a ≥ 2 fold change between two adjacent developmental stages (Figure 2A, Table S3) (ANOVA, multiple-testing corrected *P*-value < 0.01). The difference between Endo and HE is modest, consisting of only 298 DEGs, reflecting the recent endothelial origin of HE. The largest transcriptome change occurs between pre-HSCs and FL HSCs, which differentially express 3,549 genes.

**Figure 2.**
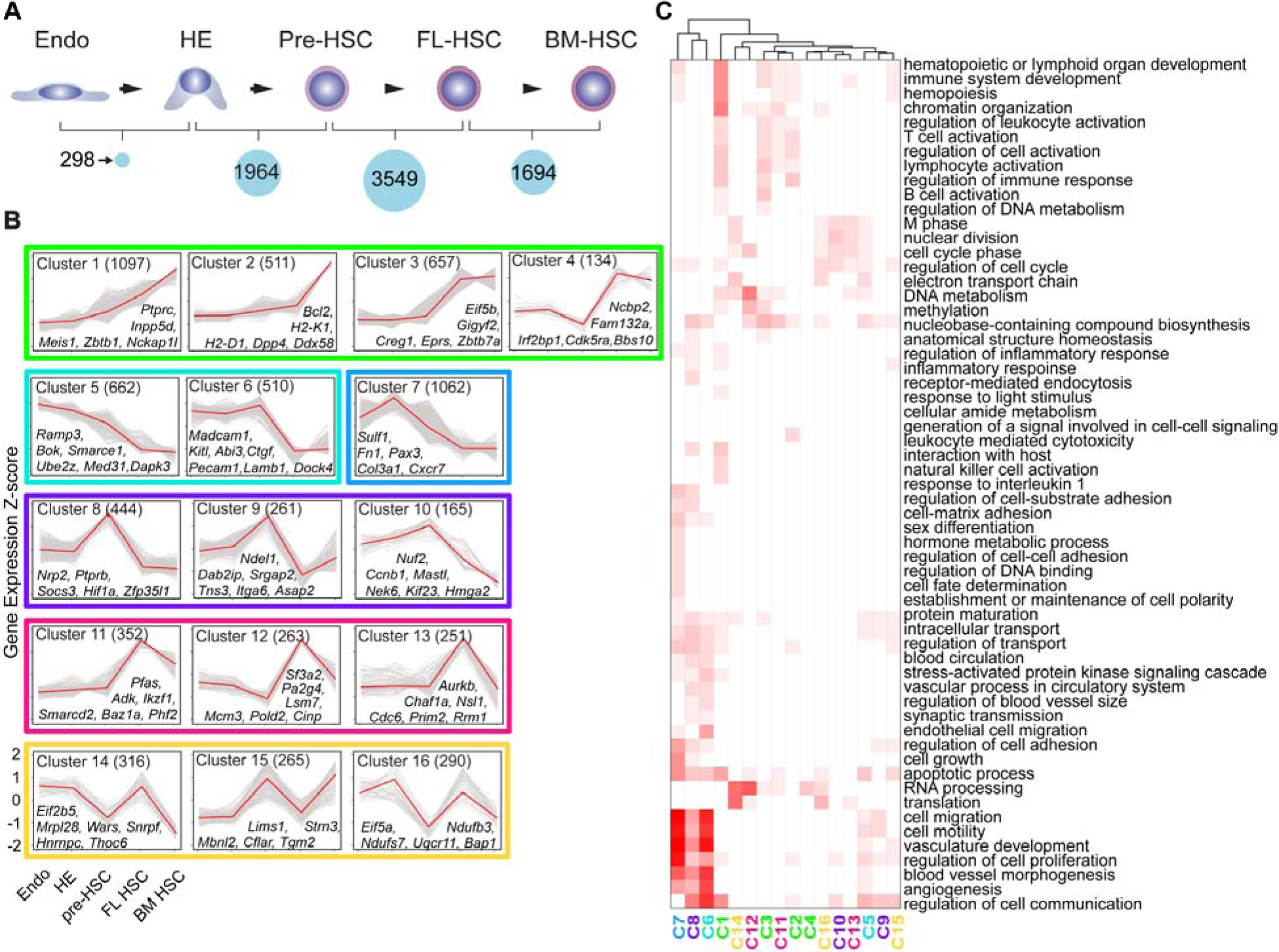
Transcriptome dynamics during HSC ontogeny. **(A)** Number of differentially expressed genes (DEGs) between adjacent developmental stages. DEGs were identified using False Discovery Rate < 0.01 and fold change ≥ 2. “Pre-HSC” includes type I pre-HSCs, type II pre-HSCs, and HSCs in intra-aterial clusters. **(B)** Sixteen expression clusters during HSC ontogeny. Expression profiles were clustered using the STEM algorithm. Y-axis values are gene-wise Z-score of FPKM values. Y and Z-axis values are indicated on the bottom left-hand graph. For each cluster, the numbers of genes are shown in parentheses. The 16 Clusters are further organized into 6 color-coded groups based on their overall expression profiles. Group 1 (clusters 1-4) are boxed in green, group 2 (clusters 5-6) in cyan, group 3 (cluster 7) in blue, group 4 (clusters 8-10) in purple, group 5 (clusters 11-13) in magenta and group 6 (clusters 14-16) in yellow. Example genes for each cluster are also shown. (**C)** Enriched Gene Ontology (GO) biological process terms among expression clusters. Color shade is proportional to the minus logarithm of enrichment *P*-value.

Using the Short Time-series Expression Miner (STEM) algorithm (Ernst et al., 2005), we identified 16 expression clusters among the 5,025 genes with ≥ 2 fold changes between two adjacent developmental stages (Figure 2B). The expression clusters are further categorized into six groups based on their expression dynamics across developmental stages. Group 1 genes (clusters 1-4) gradually increase in expression over HSC ontogeny, with peak levels in FL and/or BM HSCs. Genes in group 1 are enriched for Gene Ontology (GO) terms associated with functional HSCs, including hematopoietic or lymphoid organ development, immune system development, T cell activation, and B cell activation (Figure 2C). Group 2 genes (clusters 5-6) exhibit the opposite trend, i.e. are expressed at the highest levels in Endo cells, and are enriched for GO terms such as endothelial cell migration, cell motility, vasculature development, and angiogenesis. Group 3 (cluster 7) genes are most highly expressed in HE and are enriched for GO terms such as cell fate determination and inflammatory response terms. Group 4 genes (clusters 8-10) are most highly expressed in pre-HSCs and enriched for genes involved in inflammatory responses, cell migration, regulation of cell communication, and regulation of cell cycle. Group 5 genes (clusters 11-13) are most highly expressed in FL HSCs, and like genes in Group 1 are enriched for terms associated with functional HSCs. Genes in groups 1 and 5 are also enriched for RNA processing, DNA metabolism, and methylation terms. Group 6 genes (clusters 14-16) have oscillating expression patterns. In summary, the transcriptome analysis revealed a modest change in transcriptomes between Endo and HE and a large change accompanying the maturation of pre-HSCs to functional HSCs. Inflammation is associated with HE and pre-HSCs, consistent with functional studies demonstrating a role for inflammatory signaling in HSPC formation (Espin-Palazon et al., 2014; Li et al., 2014; Sawamiphak et al., 2014).

### Diverse activity profiles of developmental enhancers during HSC ontogeny

We developed a low-cell-number protocol to perform ChIP-Seq with as few as 20,000 cells to determine genome-wide distribution of histone modifications (Gao et al., 2019). We validated our protocol for H3K4me3 with the human lymphoblastoid cell line, GM12878. Treating ChIP-Seq peaks called using a conventional protocol as the gold standard, data generated by our protocol had an area under the ROC curve (auROC) value of 0.97, indicating excellent overall concordance (Gao et al., 2019). We profiled four histone modification marks: H3K4me1, H3K4me3, H3K27me3, H3K27ac across the five developmental stages. The combination of these four marks identifies enhancers and promoters of different activities (Calo and Wysocka, 2013). The average sequencing depth is 29 million reads per sample, which is above the sequencing depth of 20 million reads recommended by the ENCODE consortium (Landt et al., 2012). 68% of total reads could be uniquely mapped to the mouse genome (Figure S3, Table S4). The average (across 4 histone marks) genome-wide correlation of biological replicates was 0.957 (Figure S4).

Using the Chromatin Signature Identification by Artificial Neural Network (CSI-ANN) algorithm (Firpi et al., 2010) and our epigenomic data, we predicted ∼10,500 active enhancers (H3K4me1+, H3K4me3-, H3K27ac+, H3K27me3-) in each population (False Discovery Rate (FDR) < 0.05). Multiple lines of evidence corroborated our enhancer predictions. First, using ChIP-qPCR and additional purified cells, we confirmed the epigenetic signals of a select set of predicted enhancers (Figure S5). Second, our enhancer catalog is supported by additional public data, including sequence conservation across 20 mammalian genomes (34% of all predicted enhancers), a 53% overlap with reported enhancers in mouse BM HSCs (Lara-Astiaso et al., 2014), and 34% overlap with ChIP-Seq peaks of ten hematopoietic TFs (34%) (Wilson et al., 2010). Overall, 75% of our predicted enhancers were supported by at least one line of evidence (Figure 3A). The catalog also includes a number of well-known HSC enhancers identified using transgenic reporter assays in mouse, such as the *Runx1* +23 enhancer located 23.5 kb downstream of the transcription start site in exon 1 (Bee et al., 2009), the *Ly6a* (Sca1) enhancer (Ma et al., 2002), the *Tal1* (*Scl*) 3’ enhancer (Sanchez et al., 1999), the *Fli1* (Pimanda et al., 2007), *Gata2* (Lim et al., 2012), and *Erg* enhancers (Thoms et al., 2011) (Table S5).

**Figure 3.**
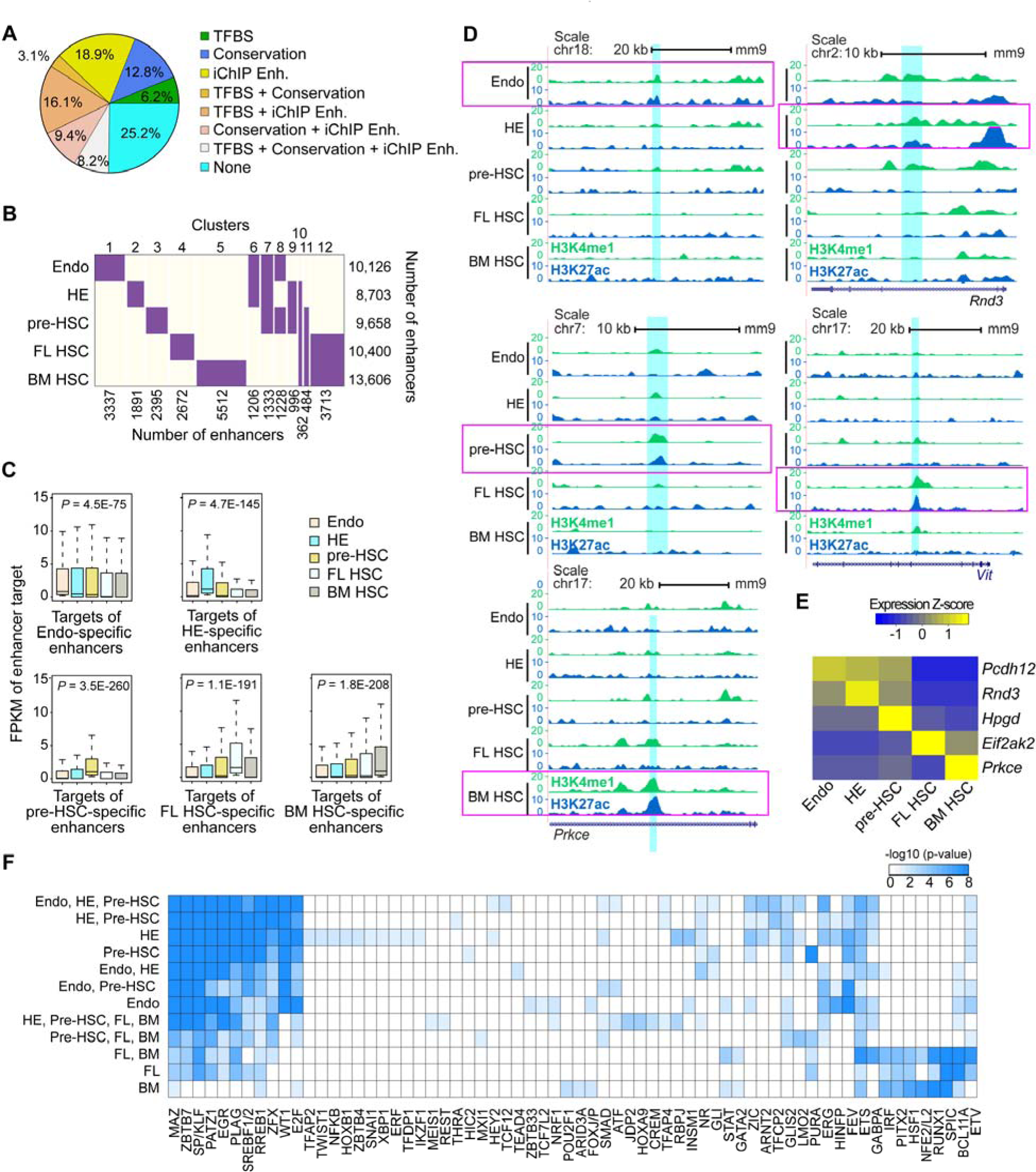
Repertoire of active enhancers during HSC ontogeny. **(A)** Supporting evidence for predicted enhancers. TFBS, transcription factor binding peaks of ChIP-Seq data from Wilson *et al*. (Wilson et al., 2010); Conservation, conserved sequence across 20 mammalian genomes; iChIP Enh, enhancers reported in Lara-Astiaso *et al*. (Lara-Astiaso et al., 2014). (**B**) Clustering of enhancer activity profile across five developmental stages. Active enhancers are predicted using the epigenetic signature and CSI-ANN algorithm. Purple, active enhancers; ivory, inactive enhancers. Numbers of enhancers in each cluster are shown to the left of the clustergram. (**C**) Boxplots of expression profile of target genes of stage-specific enhancers. P-values, differential expression between stage-specific enhancer targets and targets of the rest of the enhancers using t-test. (**D**) Genome browser view of example developmental-stage-specific enhancers (2 kb) in Endo, HE, pre-HSC, FL HSC and BM HSC. Vertical cyan boxes denote enhancers. Magenta horizontal boxes highlight tracks for each cell population. (**E**) Target gene expression patterns of developmental-stage-specific enhancers in (D). (**F**) TFs whose DNA binding motifs are enriched in stage-specific enhancers. Stage specificities are indicated on the left of each column. BM is BM HSC and FL is FL HSC. Color shade is proportional to the minus logarithm of enrichment *P*-value.

Lara-Astiaso et al. (Lara-Astiaso et al., 2014) used an epigenomics-based annotation strategy to catalogue 37,548 enhancers in cell types spanning the developmental hierarchy from adult BM HSCs to terminally differentiated hematopoietic cell types such as lymphocytes and macrophages. We queried what fraction of our enhancers (26,794 in total, excluding enhancers of Endo) overlap with the enhancers identified in that study. Strikingly, only 4,058 enhancers are shared between the two datasets. The majority of the enhancers from our study overlapping with those identified by Lara-Astiaso et al. (Lara-Astiaso et al., 2014)correspond to our BM HSC enhancers (2421 enhancers), whereas there was less overlap with HE, pre-HSC, and FL HSC enhancers (Table S6). The data comparison suggests that the enhancer repertoire for specifying HSC fate is quite distinct from that for multi-lineage differentiation from adult HSCs. Consensus clustering based on the enhancers’ activity profiles across the five developmental stages organized them into twelve clusters (Figure 3B). Clusters 1-5 represent developmental stage-specific enhancers active in Endo, HE, pre-HSCs, FL HSCs, or BM HSCs, accounting for 29.4% of all enhancers. This large percentage of stage-specific enhancers suggests the transcriptional regulatory program during HSC development is quite dynamic. In fact, we found no enhancer active across all five developmental stages. Cluster 6 represents enhancers uniquely active in endothelial cells (Endo and HE). Enhancers in cluster 7 are active in Endo, HE and pre-HSCs. Enhancers in cluster 12 are only active in functional HSCs.

Using the Integrated Method for Predicting Enhancer Targets (IM-PET) algorithm (He et al., 2014), we linked 25,541 active enhancers to their target promoters at the five developmental stages, resulting in a total of 120,257 enhancer-promoter (EP) interactions (Table S7). Expression levels of genes targeted by developmental-stage-specific enhancers are significantly higher at the corresponding stage than at other stages, supporting our EP predictions (Figure 3C). On average, 2.3 enhancers contact a promoter in all populations (Figure S6A), while 4.4 promoters contact an enhancer (Figure S6B), consistent with results from recent large-scale chromatin interaction studies of EP interactions (Kieffer-Kwon et al., 2013; Li et al., 2012; Sanyal et al., 2012). Only ∼14.3% of the E-P pairs had no intervening promoters from unrelated genes (Figure S6C). 35.0% of the predicted E-P interactions were developmental-stage-specific (Figure S6D). For example, protocadherin 12 (*Pcdh12*, also called VE-cadherin 2) encoding a member of the cadherin superfamily that plays an important role in cell-cell interactions at inter-endothelial junctions (Rampon et al., 2005; Telo et al., 1998), is predicted to be regulated by an Endo-specific enhancer. A HE-specific enhancer is illustrated for the Rho family GTPase 3 (*Rnd3*, also called *RhoE*) gene, which encodes an atypical small GTPase that induces cytoskeleton organization and cell rounding (Guasch et al., 1998; Nobes et al., 1998; Pellegrin and Mellor, 2007). A pre-HSC-specific enhancer was identified targeting prostaglandin dehydrogenase 1 (*Hpgd*) encoding the enzyme that mediates the metabolism of prostaglandin, a stimulator of HSPC formation (North et al., 2007) (Figure 3D). Examples of FL HSC and BM HSC-specific enhancers are the FL HSC-enhancer that targets eukaryotic translation initiation factor 2 alpha kinase 2 (*Eif2ak2*), a regulator of translation initiation, and the BM HSC-specific enhancer of protein kinase C epsilon (*Prkce*) that plays a role in asymmetric division and maintenance of long-term HSCs (Altman and Kong, 2016; Hazen et al., 2011; Ivanova et al., 2002). The putative target genes of the enhancers exhibit the same developmental-stage-specific expression patterns as their enhancers, consistent with regulation by the predicted enhancers (Figure 3E).

To identify TFs that occupy the enhancers, we conducted a TF motif enrichment analysis using a set of 9,500 enhancers from 18 non-hematopoietic cell types as the background. In total, we identified 78 enriched TF motif families across the 12 enhancer clusters (Figure 3F). Motifs of many known hematopoietic TFs were identified, including RUNX1, GATA2, HEY2, LMO2, and HOXA9. Motifs of broadly expressed TFs such as Kruppel like factors (KLFs) and specificity proteins (SPs) are also enriched at hematopoietic enhancers. The enriched motifs are organized into prominent groups. TF motifs on the left side of the plot, including, for example, MAZ, ZBTB7, SP/KLF, and PATZ2 are strongly enriched in the enhancers active in endothelial-like cells (E, HE, pre-HSC), become less enriched in FL HSC enhancers, and are not enriched in enhancers active in only BM HSCs. Most of the TFs that recognize these motifs are not thought of as canonical HSC TFs, but may be important during the process of pre-HSC formation from endothelial cells. The opposite pattern is observed for TF motifs on the right side of the plot, which are more canonical HSC TF motifs that are not enriched in the enhancers active in E-like populations. A small group of TF motifs are enriched only in active HE enhancers. Interesting, two of the implicated HE-specific TFs, SNAI1 and TWIST1, are involved in endothelial to mesenchymal transitions (EndMT), a process similar to EHT (Hulshoff et al., 2019).

### Repertoire of epigenetically primed enhancers during HSC ontogeny

Both active and epigenetically primed enhancers are marked by H3K4me1, but can be distinguished by the presence (active) or absence (primed) of H3K27ac, H3K27me3, and H3K4me3 modifications (Calo and Wysocka, 2013). Primed enhancers play important roles in cell fate transitions during development and differentiation in response to environmental cues (Wang et al., 2016), in maintaining epigenetic memory at cis-regulatory elements (Bevington et al., 2017), and for rapid environmental responses (Gosselin et al., 2017). Using the CSI-ANN algorithm (Firpi et al., 2010) and our epigenomic data, we predicted between 2574 and 3342 primed enhancers across the five developmental stages (Figure 4A, Table S8). The number of active enhancers is approximately 3.5 times greater than that of primed enhancers at all developmental stages. Enhancers can be activated with or without prior epigenetic priming; activation without priming is considered *de novo* activation. The two different activation mechanisms suggest different developmental functions of the enhancers. We found that the vast majority of hematopoietic enhancers are activated *de novo* without priming (Figure 4B). The target genes of enhancers that are primed before activation tend to have high expression fold changes at the subsequent developmental stage (Figure S7).

**Figure 4.**
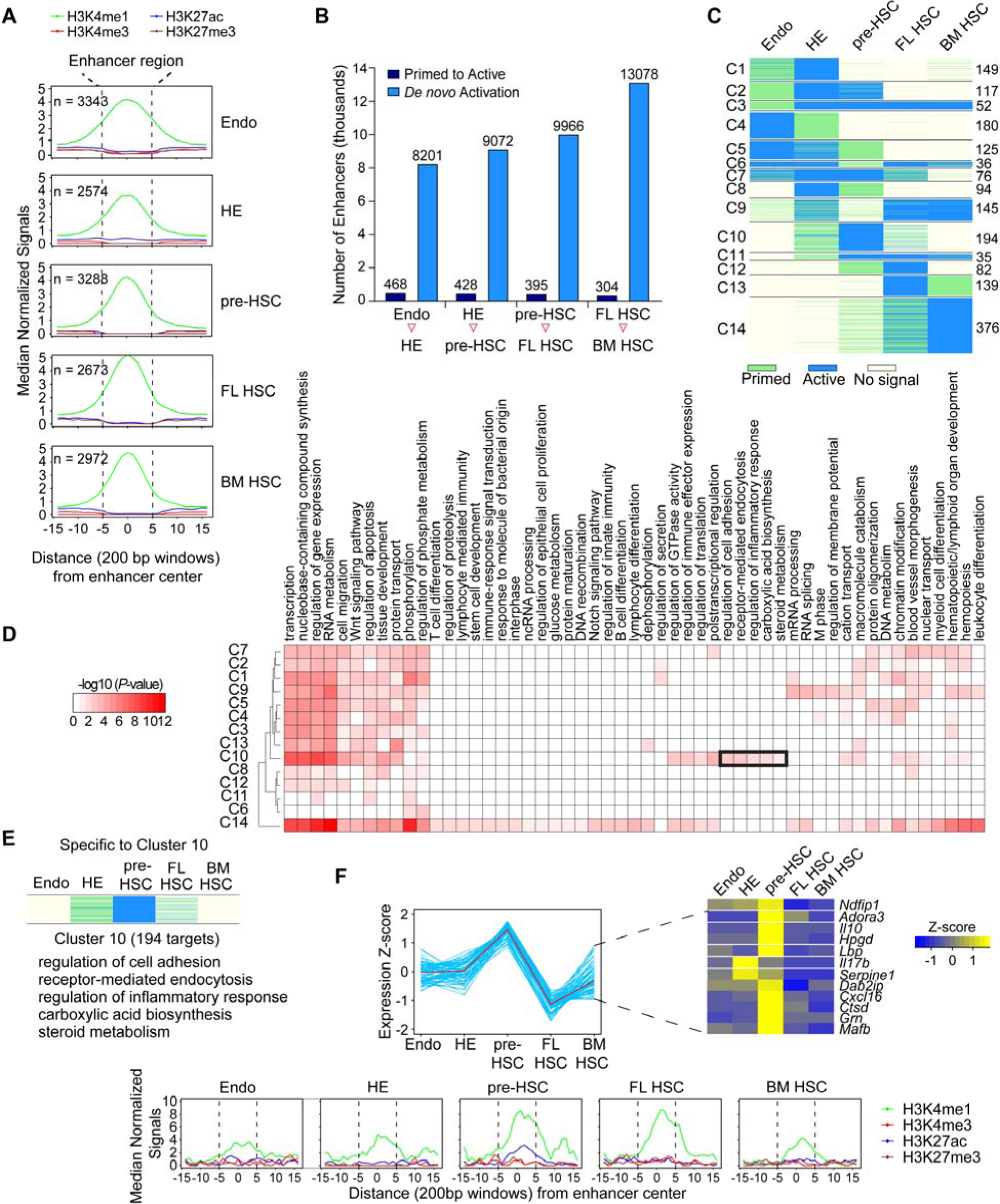
Primed enhancers during HSC ontogeny. **(A)** Metagene plot of normalized histone mark ChIP-Seq signals at predicted primed enhancers (n = total number of primed enhancers identified in each stage) **(B)** Dynamics of primed and *de novo* activated enhancers during HSC ontogeny between each adjacent stage. **(C)** Clustering based on activation pattern of the primed enhancers during HSC ontogeny. Cluster number is on the left and the number of enhancers within each cluster on the right. (**D**) Enriched GO terms of genes targeted by primed enhancers within each cluster. Black rectangle highlights C10-specific GO terms specific for pre-HSCs. (**E)** Expression profile and enriched GO terms of predicted target genes of enhancers in cluster 10. (**F**) Top, expression of inflammatory response genes in cluster 10. Bottom, metagene plot of normalized histone mark ChIP-Seq signals at primed enhancers targeting the inflammatory response genes in cluster 10.

We next investigated the activation pattern of the primed enhancers during HSC ontogeny (Figure 4C). We first identified primed enhancers that were activated either in the previous or subsequent developmental stage, and clustered these enhancers using their epigenetic profiles across the five stages. We observed diverse priming-and-activation patterns that can be grouped into 14 distinct clusters (Figure 4C, Table S9). Several groups consist of enhancers with a strong pattern of priming followed by activation (clusters C2, C3, and C12), or conversely, activated then primed at a subsequent stage (clusters C4 and C8). We also observed enhancers that gradually became primed or activated over several developmental stages (clusters C5 and C14). The different groups of enhancers regulate distinct gene pathways (Figure 4D). Cluster C10 enhancers (n = 194) and their predicted target genes are of particular interest; they are primed in HE, become active in pre-HSCs, gradually lose their H3K27ac marks and revert to primed enhancers in FL HSCs, and subsequently lose their H3K4me1 marks and are deactivated in BM HSCs. The target genes within cluster C10, which are likely involved in regulating the expression of genes important for the specification of HE and formation of pre-HSCs, are enriched for several unique pathways, including regulation of cell adhesion, receptor-mediated endocytosis, regulation of inflammatory response, carboxylic acid biosynthesis and steroid metabolism (Figure 4E). Several of these pathways, including inflammatory signaling, cholesterol biosynthesis, and vitamin D synthesis (i.e. steroid metabolism) were shown to promote the formation of HSPCs from HE (Cortes et al., 2016; Espin-Palazon et al., 2014; Gu et al., 2019; Li et al., 2014; Sawamiphak et al., 2014). Consistent with the enhancer activation pattern, expression of the target pathways is also highest in pre-HSCs (Figure 4F, left panel). For instance, expression of the inflammatory response genes is highest in pre-HSC (Figure 4F, right panel), these genes’ enhancers are primed in HE, and become active in pre-HSC (Figure 4F, bottom panel). In summary, our data suggest genes in multiple pathways are regulated by primed enhancers that respond to signaling pathways important for pre-HSC formation (Figure 4E).

### Core transcriptional regulatory circuitries underlying developmental transitions

Little is known about the transcriptional regulatory networks (TRNs) during HSC formation from arterial endothelium. We recently developed the Target Inference via Physical Connection (TIPC) algorithm for inferring condition-specific TRNs by integrating RNA-Seq and histone mark ChIP-Seq data (Gao et al., 2019). TIPC first computes probability scores for three key components of transcriptional regulation, including probability of a DNA sequence being an enhancer, probability of a TF binding to an enhancer based on the TF motif model and the enhancer sequence, and probability of enhancer-promoter interaction. The overall score for a TF regulating a target gene is the product of the three component probabilities and the expression level of the TF (Figure 5A). Using a set of gold standard TF-target gene pairs in mouse embryonic stem cells, we demonstrated that TIPC achieves improved accuracy compared to several state-of-the-art methods (Gao et al., 2019). To increase robustness, we first scanned the enhancer sequences for each TF motif using the Find Individual Motif Occurrence (FIMO) software (Grant et al., 2011) (*P-*value < 1e-5). We then applied TIPC to construct condition-specific TRNs for the five developmental stages. On average, each TRN consists of 381 TFs and 6,530 target genes.

**Figure 5.**
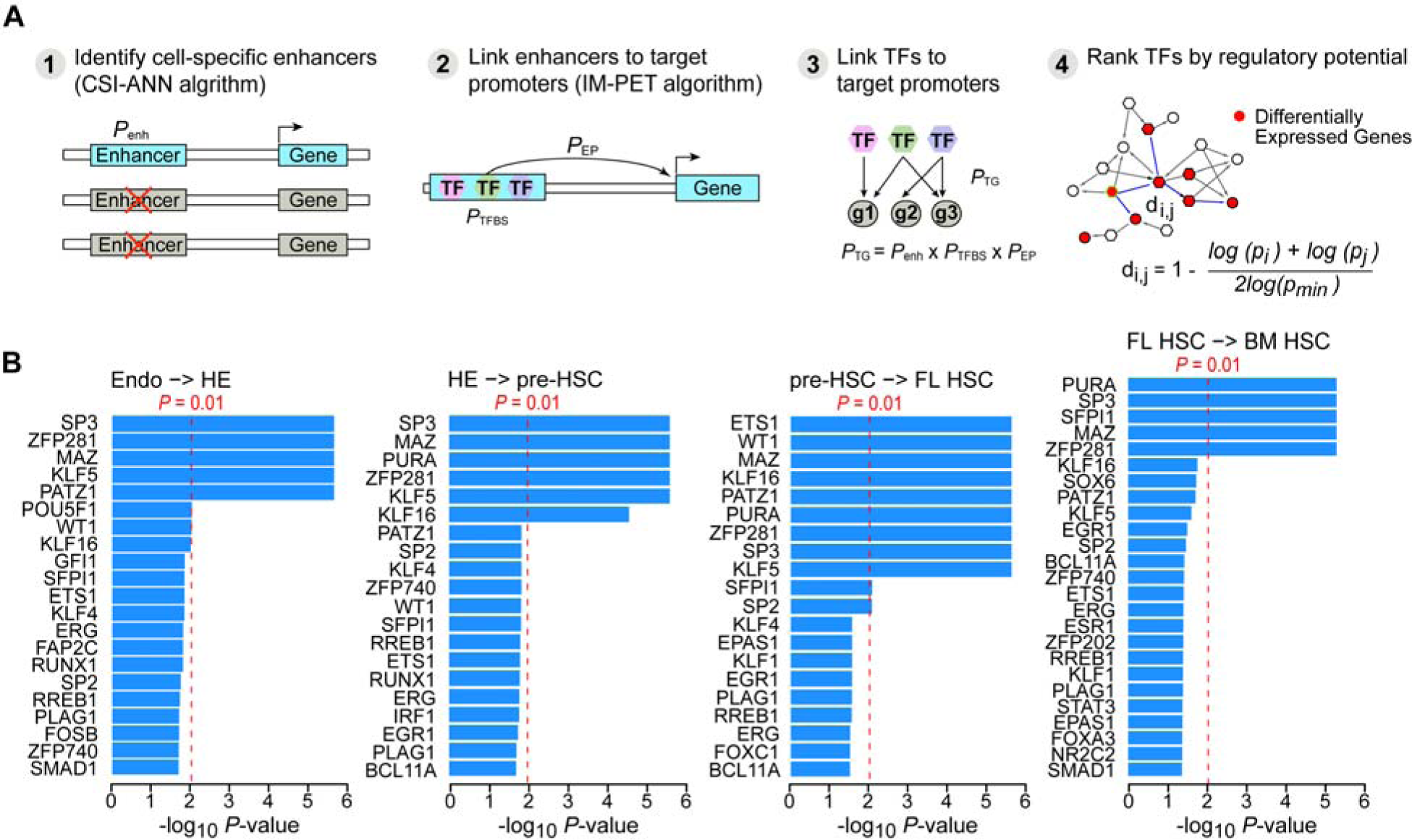
Developmental-stage-specific transcriptional regulatory networks (TRNs) and key regulators in the networks. **(A)** Schematic for the computational framework for constructing the TRNs. The framework consists of four key steps: 1) identifying stage-specific enhancers using epigenetic signatures; 2) linking enhancers to target promoters using the IM-PET algorithm; 3) linking TFs to target promoters using a probabilistic scoring function *p_TG_*; 4) identifying key TFs based on their regulatory potential on differentially expressed genes. (**B)** Key TFs predicted for four developmental transitions.

To identify key TFs regulating each developmental transition in HSC ontogeny, we first merged developmentally adjacent TRN pairs and represented each gene in the merged network based on its level of differential expression during the transition. Next, we identified key TFs in the differential TRN using a network propagation-based method (Gao et al., 2019). The rationale for our method is that a key TF likely exerts its influence (either directly or indirectly) on the entire set of differentially expressed genes via a shorter regulatory path compared to paths from less important TFs. We predicted the sets of key TFs for each developmental transition using a heuristic similar to described in Mogrify (Rackham et al., 2016) (Figure S8, 5B, Table S10). Our algorithm predicted known regulators of HSC ontology, such as RUNX1 and GFI1, which are essential for EHT (Lancrin et al., 2012; North et al., 1999). Two other TFs predicted by our algorithm, FOSB and SPI1, were sufficient to reprogram endothelial cells to HSCs when transiently expressed with RUNX1 and GFI1 (Lis et al., 2017; Sandler et al., 2014). Additional validated TFs that play a role in HE and/or HSPC formation predicted by our algorithm were SP3, ETS1, ERG, and TFs belonging to the BMP-SMAD1 pathway (Gilmour et al., 2019; Park et al., 2018; Sugimura et al., 2017; Zhang et al., 2014). Our algorithm also predicted TFs that have yet to be implicated in HSC formation, such as Myc-associated zinc finger (MAZ)

### Novel roles of SP3 and MAZ in embryonic hematopoiesis

Two of the most highly ranked TFs predicted by our algorithm to be involved in multiple stages of HSC ontogeny were MAZ and SP3. No role for MAZ in any aspect of hematopoiesis has been reported, although MAZ is a regulator of inflammation and Notch signaling in other tissues (Liu et al., 2016; Ray et al., 2004). Germline deletion of *Sp3* was shown to decrease hematopoiesis in the mouse fetus and the lymphoid output of FL HSCs upon transplantation into adult recipient mice (Van Loo et al., 2003), and deletion of *Sp3* in ES cells decreased the number of primitive erythroid progenitors generated in differentiation cultures (Gilmour et al., 2019). However, in neither study of *Sp3* were the associated defects examined or implicated at the level of HE. To validate the predictive power of our algorithm, we examined the role of SP3 and MAZ in HSPC formation by using CRISPR-Cas9 to knock out the orthologs of *Sp3* and *Maz* in zebrafish embryos (Table S11, 12). Both genes have two orthologs in zebrafish (*sp3a* and *sp3b* for *Sp3*, *maza* and *si:ch211-166g5.4* for *Maz*). We knocked out both zebrafish orthologs of *Sp3* or *Maz* and examined *runx1* and *c-myb* expression in the dorsal aorta at 27 and 30 hours post fertilization (hpf), respectively, by whole-mount *in situ* hybridization (WISH) (Figure 6A-B) (Bertrand et al., 2010; Bertrand et al., 2008; Lam et al., 2009; Lam et al., 2010). Knockout of *sp3a* and *sp3b* decreased *runx1* and *c-myb* expression in the dorsal aorta (Figure 6C, D), as did knockout of the two *Maz* orthologs (Figure 6E, F). The percentage of kdrl:mCherry^+^ myb:GFP^+^ HSPCs was also significantly reduced in both *Sp3* knockout and *Maz* knockout zebrafish embryos compared to wild type embryos (Figure 6G, H). To exclude the possibility that the reductions in HE and HSPCs were due to antecedent vascular abnormalities, we examined the integrity of blood vessels at 48 hpf in *Tg(kdrl:mCherry)* transgenic embryos (Tamplin et al., 2015). There were no significant differences in the vascular sprouting patterns, the function of intersegmental blood vessels (ISVs), or in the diameters of the ISVs, dorsal aortae, or posterior cardinal veins between wild type and knockout embryos of either genotypes, indicating the decreases in *runx1* and *myb* expression, and the reduced frequencies of HSPCs in *Sp3* and *Maz* knockout embryos were specific to HE (Figure S9).

**Figure 6.**
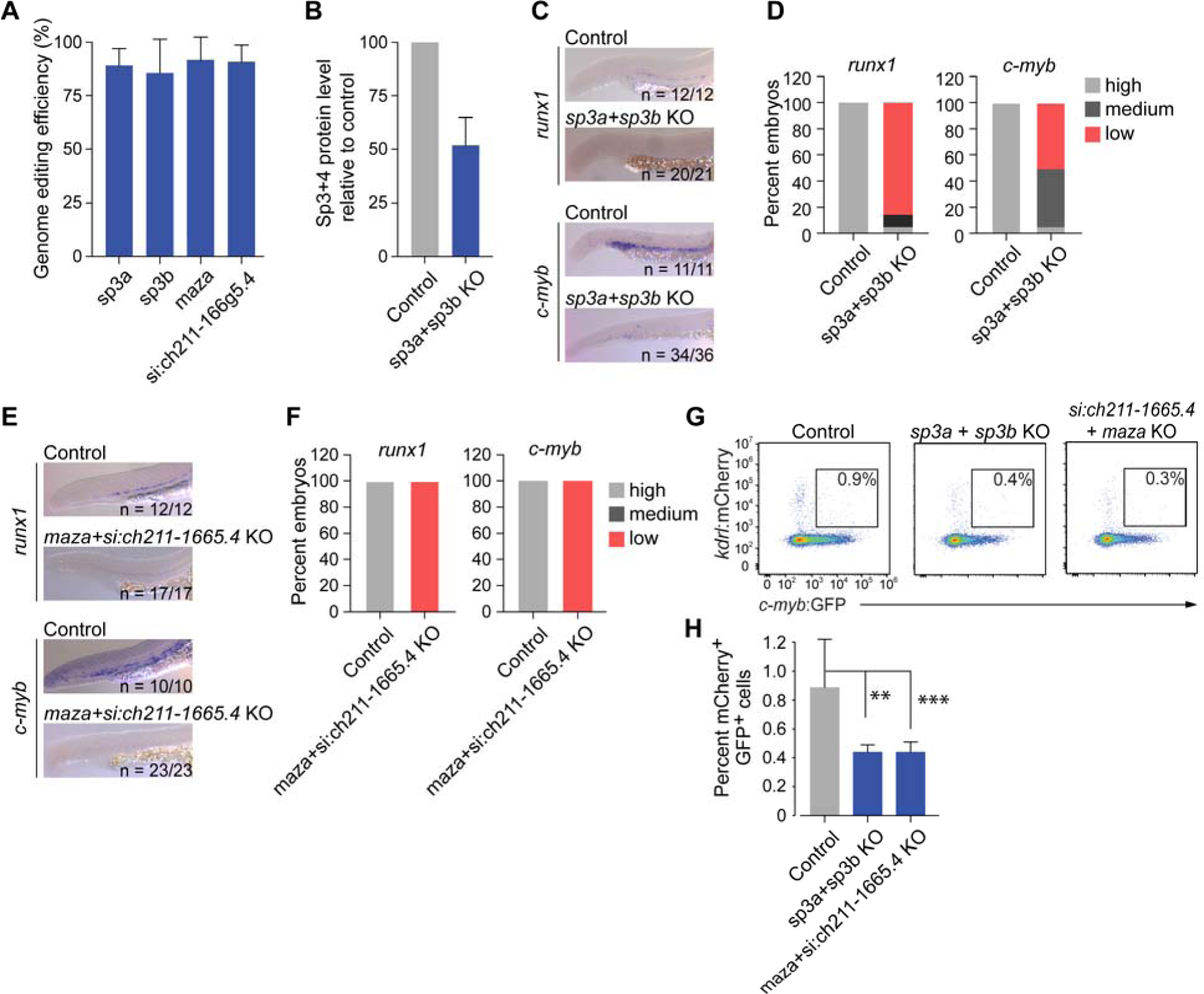
*Sp3* and *Maz* orthologs promote HSPC formation in zebrafish embryos. **(A)** Genome editing efficiency of each sgRNA determined using the Tracking of Indels by DEcomposition assay (TIDE). Data represent mean ± SD of three replicate experiments. **(B)** Relative protein level of SP3 and SP4 in the knockout (KO) embryos. Data represent mean ± SD, n = 9. **(C)** Representative whole-mount *in situ* hybridization (WISH) images for *runx1* and *c-myb* in the AGM region of 27 hpf embryos. Embryos were injected with CRISPR guide RNAs targeting the *Sp3* orthologs *sp3a* and *sp3b*. The numbers of embryos with reduced *runx1* or *c-myb* expression relative to the number examined are indicated. (**D**) Percentage of embryos with high, medium, and low *runx1* and *c-myb* expression following injection of sgRNAs targeting *Sp3* orthologs, and controls. (**E**) Representative WISH images for *runx1* and *c-myb* in the AGM region of 30 hpf control and KO embryos injected with CRISPR guide RNAs targeting both *Maz* orthologs, *maza* and *si:ch211-166g5.4*. (**F**) Percentage of embryos with high, medium, and low reporter gene expression after injection of sgRNAs targeting the *Maz* orthologs. (**G**) Representative flow cytometry showing gating strategy for mCherry^+^ GFP^+^ HSPCs from embryos transgenic for flk1:mCherry and c-myb:GFP reporter genes following KO of Sp3 or Maz orthologues. **(H)** Frequency of mCherry^+^ GFP^+^ HSPCs in wild type, *Sp3*, and *Maz* knockout embryos (KO). *** *P* ≤ 0.01, ** *P* ≤ 0.05. Student’s t-test.

To further evaluate if SP3 loss affected the specification of HE, we enumerated IACs and HE cells in E10.5 mouse embryos, during the peak of EHT, by whole mount confocal microscopy (Yokomizo et al., 2012). Loss of SP3 reduced the number of CD31^+^ RUNX1^+^ c-Kit^+^ IACs and CD31^+^ RUNX1^+^ c-Kit^-^ HE cells in the dorsal aorta, consistent with a role for SP3 in specifying HE (Figure 7A-C). To examine the impact of SP3 loss on functional HE cells, we purified arterial endothelial cells (CD31^+^ CD144^+^ ESAM^+^ CD44^+^ CD41^lo/-^) from the dorsal aorta of E10.5 wild type and *Sp3* deficient embryos and determined the frequency of HE cells capable of producing blood cells in a limiting dilution assay (Figure 7D, E). The frequency of functional HE cells in the sorted arterial endothelial cell populations was reduced by 75% in *Sp3^-/-^* embryos, indicating that SP3 promotes the specification of HE cells in the dorsal aorta of mouse embryos. Interestingly, loss of SP3 had no effect on colony formation from the yolk sac (Figure 7F), suggesting that SP3 does not regulate the specification of yolk sac HE. This further validates the ability of our algorithm to predict TFs required for definitive hematopoiesis in the major arteries of the embryo.

**Figure 7.**
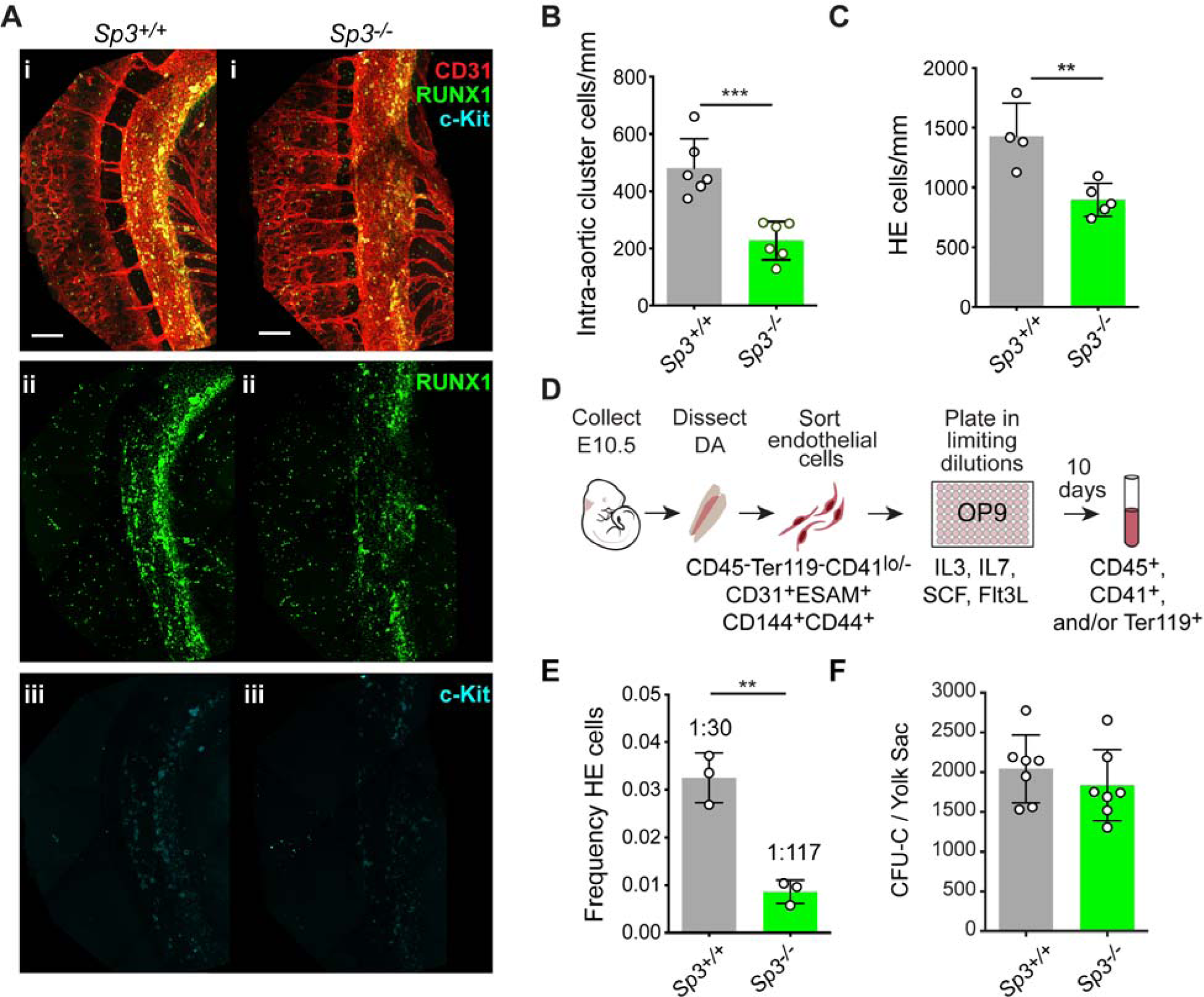
SP3 regulates the frequency of HE cells in the dorsal aorta of mouse embryos. (**A**) Confocal z-projections (z-intervals = 2.5 µm) of dorsal aortas from E10.5 *Sp3^+/+^* and *Sp3^-/-^* mouse embryos. Embryos were immunostained for CD31 (i), RUNX1 (i,ii) and c-Kit (i,iii). Scale bars: 100 µm. (**B**) Quantification of CD31^+^RUNX1^+^c-Kit^+^ intra-aortic hematopoietic cluster cells in the dorsal aortas of E10.5 embryos (mean ± SD, unpaired two-tailed t-test). (**C**) Quantification of HE cells (CD31^+^RUNX1^+^c-Kit^-^) in the dorsal aorta (mean ± SD, unpaired two-tailed t-test). (**D**) Illustration of a limiting dilution assay to determine the frequency of functional HE cells in a purified population of arterial endothelial cells. **(E)** Frequency of HE cells in CD45^-^ Ter119^-^CD31^+^ESAM^+^CD144^+^CD44^+^CD41^lo/-^ cells purified from the dorsal aortas of E10.5 *Sp3^+/+^* or *Sp3^-/-^* embryos (mean ± SD, unpaired two-tailed t-test). Each replicate consisted of pooled cells from separate litters of E10.5 embryos collected in independent experiments. *** *P* ≤ 0.001, ** *P* ≤ 0.01. (**F**) CFU-Cs per E10.5 YS (mean ± SD). Data are from three independent litters.

### Discussion

Here we provide the first comprehensive map of the transcriptome and the epigenome throughout HSC ontology from the major arteries. We defined transcriptional enhancers that are primed or activated at five distinct stages of endogenous definitive hematopoiesis in the AGM. Thirty percent of the identified enhancers are specifically active at one developmental stage, which exceeds the percentage of differentially expressed genes (12%) between any two adjacent developmental stages. These results suggest that the cis-regulatory landscape is more dynamic than gene expression during HSC ontogeny, and that enhancer patterns may be finer-grained descriptors of cellular differentiation stages than gene expression. It is possible that the excess of stage-specific enhancers relative to stage-specific gene expression may be driven by a need for combinatorial regulation by multiple enhancers to achieve specific complex expression patterns. Alternatively, or in addition, an inherent redundancy may be built into the regulatory program to achieve evolutionary robustness (Long et al., 2016), a phenomenon also reported in erythropoiesis (Xu et al., 2012), forebrain development (Visel et al., 2013), and embryogenesis in fruit fly (Cannavo et al., 2017).

Our study also confirms the importance of the broadly expressed TF SP3 in HSC ontogeny (Gilmour et al., 2014), and identifies a role for the related protein MAZ. SP3 and MAZ are zinc finger proteins that recognize similar, although distinct GC-rich DNA sequences (Song et al., 2001; van Loo et al., 2007). Twenty seven percent of human promoters contain GC-rich regions, making GC boxes the most prevalent promoter motifs (FitzGerald et al., 2004). The ubiquitously expressed TFs that bind GC boxes are capable of regulating a large number of genes involved in fundamental cellular processes including proliferation, apoptosis, migration, adhesion, and angiogenesis. The tissue specific activity of these TFs is determined by their interaction with lineage specific TFs. For example, SP3 was reported to bind regions occupied by RUNX1 and PU.1 in myeloid cells (Lennartsson et al., 2003). We determined, using publicly available ChIP-seq data from differentiating murine hematopoietic cells (Lichtinger et al., 2012), that RUNX1 binds at a significant number of enhancers and promoters of predicted SP3 target genes in HE (hypergeometric test, *P*-value < 0.05), suggesting that RUNX1 and SP3 may be cooperating to regulate these genes. Interestingly, we see that at later stages of HSPC ontology, SP3 appears to be cooperating with different hematopoietic TFs, such as FLI1, LMO2, SCL, and GFI1/B, as determined by the significant overlap of binding of these TFs at enhancers and promoters of predicted SP3 target genes (hypergeometric test, *P*-value < 0.05) (Goode et al., 2016). Supporting this, the Sp-family protein SP1 has also been reported to physically interact with RUNX1 (Wei et al., 2008), GATA2 (Maeda et al., 2010), and the SCL complex consisting of SCL, LMO2, and GATA1/2 (Lecuyer et al., 2002).

The cell type-specific activity of broadly expressed TFs could be conferred by the chromatin landscape established by lineage-specific TFs. This model is supported by data showing that loss of SP1 in ES cells had little effect on chromatin structure, and that SP1 and SP3 primarily bind in regions of accessible chromatin marked by the presence of H3K27Ac and H3K4me3, and the absence of H3K9me3 (Gilmour et al., 2014). Our data fits with the model proposed by Gilmour and colleagues suggesting ubiquitous factors, such as SP3 and MAZ, are important for establishing a stable promoter structure that interacts with distal regulatory elements. In this model, lineage-specific factors are responsible for making the correct distal regulatory elements accessible for ubiquitously expressed factors to interact with.

The stage-specific enhancers and core transcriptional regulatory circuitries identified in this study will be an important resource for isolating transitional cell populations during hematopoietic ontogeny, as well as for the *de novo* generation of HSCs from induced pluripotent stem cells (iPSCs) or other cell sources. Currently, the majority of protocols for deriving HSCs from iPSCs recapitulate yolk sac hematopoiesis which produces a different complement of HSPCs than the major arteries (Wahlster and Daley, 2016). Protocols for *ex vivo* HSPC production could be improved by a more comprehensive understanding of endogenous definitive hematopoiesis in the major arteries, and of the differences between HE in both sites. In addition, previous attempts to reprogram endothelial cells to become HSCs utilized TFs specifically expressed in HSCs and not in endothelial cells (Lis et al., 2017; Sandler et al., 2014). The efficiency of this process could potentially be improved by also introducing TFs normally expressed during HSC ontogeny predicted to be involved in the Endo→HE, and the HE→pre-HSC transitions.

### Data availability

All data generated in this study has been deposited in the Gene Expression Omnibus (GEO) database under the accession number GSE135601.

## Acknowledgements

We thank the Research Information Services at the Children’s Hospital of Philadelphia for providing computing support, and Andrea Stout and Jasmine Zhao at the University of Pennsylvania Cell and Developmental Biology Microscopy Core facility for microscopy assistance. This work was supported by National Institutes of Health grants GM104369, GM108716, HG006130 (to K.T.), HL091724 (to N.A.S.), HD089245 (to K.T. and N.A.S.) and HD083185 (E.D.H.).

## Materials and Methods

### Mouse strains

Hemogenic and non-hemogenic endothelial cells were purified from *Runx1^tm4Dow^* embryos (Lorsbach et al., 2004). Pre-HSCs and embryonic HSCs were isolated from B6.Cg-Tg(Ly6a-GFP)G5Dzk/J embryos. Female B6C3F1/J mice were mated with male B6129SF1/J mice to generate E14.5 fetuses for purifying fetal liver HSCs. Pairs of 6-8 week old female and male B6C3F1/J and B6129SF1/J mice were used to isolate bone marrow HSCs. SP3 mutant mice (B6.129P2-Sp3^tm1Sus^/Cnrm) were from the EMMA repository (EM:02429) and genotyped as described in Kruger et al. (Kruger et al., 2007). The University of Iowa and University of Pennsylvania Offices of the Institutional Animal Care and Use Committee (IACUC) review boards approved these studies. This study was performed in accordance with the recommendations in the Guide for the Care and Use of Laboratory Animals of the National Institutes of Health. All of the animals were handled according to approved IACUC protocols.

### Zebrafish maintenance

Zebrafish were maintained according to the IACUC-approved protocols at Children’s Hospital of Philadelphia. Zebrafish were bred, and embryos were collected for microinjection within 45 min post fertilization. Post microinjection, embryos were raised in E3 medium in 100 mm Petri dishes at a density of fewer than 100 embryos per dish. All larvae were incubated at 25 °C.

### Tissue dissection and cell purification using fluorescence-activated cell sorting

#### Hemogenic and non-hemogenic endothelial cells

Embryos were collected from B6C3F1 female mice mated to *Runx1^tm4Dow^* male mice and dissected in 1× PBS + 10% FBS + 1% Penicillin-Streptomycin (Gibco). AGM regions, umbilical and vitelline vessels were separately collected and digested in 0.125% collagenase (Sigma) for 1 hour at 37 °C. Cells were washed and stained in an antibody cocktail against CD41, CD31, c-Kit, CD45 (eBiosciences) and 7AAD or DAPI (Invitrogen). Cells were sorted on a Becton Dickinson (BD) FACS Jazz (BD Biosciences) or BD Influx into 1× PBS + 20% FBS + 25 mM HEPES in a 1.5 mL LoBind tube.

#### Pre-HSCs

5 IU of pregnant mare serum gonadotropin (Sigma) and 5 IU of human chorionic gonadotropin (Sigma) were injected into 3 week old B6C3F1/J female mice by intraperitoneal injection to stimulate superovulation, and mated to (Ly6a:GFP) male mice [B6.Cg-Tg(Ly6a-GFP)G5Dzk/J] (Ma et al., 2002). AGM, vitelline and umbilical arteries were dissected from E11.5 Ly6a:GFP^+^ embryos. Pooled tissues were dissociated with collagenase (Sigma). Cells were washed and resuspended in 1× PBS + 10% FBS (Gibco) and stained with antibodies against CD31, c-Kit, ESAM and CD144. Dead cells were excluded by DAPI staining (Molecular Probes). Cells were sorted on a BD Influx into 1 mL 1× IMDM + 50% FBS in a 1.5 mL LoBind tube.

#### E14.5 FL HSCs

B6C3F1/J female mice were mated with B6129SF1/J male mice, and fetal livers dissected from E14.5 embryos. Single cell suspensions were prepared by mechanical dissociation and expelling the cells through 40 μ using ACK lysing buffer (Lonza). Cells were washed and stained with an antibody cocktail of CD150, CD117 (c-Kit), Ly-6A/E (Sca1), CD48, Ly6G/Ly-6C (Gr-1), CD11b, TER-119, CD4, CD8a, CD45R/B220, CD3ε and CD11c. Washed cells were filtered through 40 µm cell strainer twice and Hoechest 33258 stock solution was added just prior to FACS. Cells were sorted by yield sort first followed by purity sort to obtain long-term HSCs (Lin^-^ Sca1^+^c-Kit^+^ CD150^+^ CD48^-^).

#### Bone marrow HSCs

Cells were suspended and incubated with anti-CD16/32, anti-CD32 and anti-CD117 microbeads (Miltenyi Biotec). Cells were washed twice with 1× PBS + 2% FBS and fed to autoMACS (Miltenyi Biotec) to enrich for CD117^+^ cells. Enriched cells were stained with the antibody cocktail against the following antigens: CD150, CD117 (c-Kit), Ly-6A/E (Sca1), CD135, CD48, Ly6G/Ly-6C (Gr-1), CD11b, TER-119, CD4, CD8a, CD45R/B220, CD3ε and CD11c (Supplemental Table 13) at 4 °C for 15 min in the dark. Washed cells were filtered twice through 40 µm cell strainer and Hoechest 33258 stock solution was added just prior to sorting. Cells were sorted by yield sort first followed by purity sort to obtain long-term BM HSCs (Lin^-^ Sca1^+^c-Kit^+^CD150^+^CD48^-^CD135^-^).

#### E10.5 endothelial cells for limiting dilution assay

Embryos were collected from *Sp3*^+/-^ females mated to *Sp3*^+/-^ males. The yolk sac was removed for genotyping, and the embryo dissected in 1× PBS + 10% FBS (Gibco) + 1% Penicillin-Streptomycin (Gibco). The dorsal aorta was dissected out and dissociated in 0.125% collagenase (Sigma) for 40 min. Cells were washed and stained with the following antibodies: CD45-eFluor 450 (eBioscience, 48-0451-82), Ter119-eFluor 450 (eBioscience, 48-5921-82), ESAM-FITC (Biolegend, 136205), CD41-PerCP-eFluor 710 (eBioscience, 46-0411-82), VEC-PE (eBioscience,12-1441-82), CD31-PE-Cy7 (eBioscience, 25-0311-82), CD44-APC-Cy7 (BD Pharmingen, 560568), and DAPI (Invitrogen, D3571) for viability. The cells were pooled by genotype and sorted on a BD Influx into 1 mL αMEM (Thermo Scientific) containing 10% FBS and antibiotics in 1.5 LowBind Tubes.

### Limiting dilution assay

OP9 stromal cells were cultured in αMEM (Thermo Scientific) containing 10% FBS (Gibco) and 1% Penicillin-Streptomycin (Gibco). One day prior to the initiation of the HE assay the OP9 cells were plated at a concentration of 4000 cells/well in 96-well plates. Hemogenic endothelial cell limiting dilution assays were performed by culturing sorted populations on OP9 cells supplemented with 20% FBS + 1% Penicillin-Streptomycin and 10 ng/mL each of SCF, IL-3, Flt3L and IL-7 (all cytokines from PeproTech). Six dilutions were performed with 3 replicates of each. On days 7-10, wells were inspected for hematopoietic colonies using a light microscope. Samples were analyzed using a BD LSRII flow cytometer using the following antibodies: CD45-PE-Cy7 (eBioscience, 25-0451-82), c-Kit-PerCP-eFluor 710 (eBioscience, 46-1171-82), Ter119-eFluor 450 (eBioscience, 48-5921-82), CD41-FITC (Biolegend, 133904), and Aqua LIVE/DEAD (Invitrogen, L34966) for viability. Positive wells were identified as wells containing CD41^+^, CD45^+^ and/or Ter119^+^ cells. The HE frequency was calculated using ELDA software (Hu and Smyth, 2009).

### Colony forming progenitor assay

Colony forming progenitor assays were performed by culturing populations in M3434 (StemCell Technologies). Sorted populations were plated at 3.6 embryonic equivalents. Unsorted yolk sac cells were used as a positive control, and cultured at 0.2 embryonic equivalents. For SP3 colony forming assays, whole yolk sac was dissociated in 0.125% collagenase (Sigma) and 0.03 embryonic equivalents of yolk sacs were plated in replicates. Colonies were scored on a light microscope seven days after plating.

### Low-Cell-Number ChIP-Seq

We developed a sensitive protocol that requires only 20,000 cells for histone ChIP-seq (Gao et al., 2019). Briefly, purified cells were pooled into batches of 40,000 cells and crosslinked in 1% formaldehyde (Thermo Scientific) in 1× fixing buffer for 5 min at room temperature according to the vendor’s manual (Covaris). Crosslinked cells were then resuspended in 1× shearing buffer and sonicated with Covaris E220 for 780 sec using the following settings: duty factor, 5%; peak incident power, 105; cycles per burst, 200. 5% of sheared chromatin was used as the input and the remaining chromatin was divided into two equal aliquots for immunoprecipitation (20,000 cells per IP). IPs were performed using the ChIP-IT high sensitivity kit (Active Motif). IP and input samples were treated with RNase A followed by proteinase K. Cross-linking was reversed by incubation overnight at 65 °C and DNA was purified using a MinElute PCR purification kit (Qiagen). All IP DNA and 1 ng of input DNA were used for library preparation using the ThruPLEX-FD Prep Kit or ThruPLEX DNA-Seq (Rubicon Genomics). 13 and 9 cycles were used for IP DNA and input DNA, respectively at step 5. Libraries were sequenced on Illumina HiSeq 2500 sequencer in single-end mode with read length of 50 nt.

### ChIP-qPCR

After the ChIP step in the Low-Cell-Number ChIP-Seq protocol, purified DNA from matched IP sample and input sample were subjected to qPCR analysis using iQ SYBR Green Supermix. The fold enrichment was computed as 2^-ΔΔCt^, where ΔΔCt is the difference in ΔCt values between Ct is the cycle number difference between a target region (well-known or predicted enhancer) and a negative control region (without any histone modification signal). Primer sequences are included in Table S14.

### ChIP-Seq data processing

Sequencing reads were mapped to the mouse genome (mm9) using Bowtie2 (v2.2.2) with default parameter setting (Langmead et al., 2009). Only uniquely mapped reads with fewer than 2 mismatches were used to compute a normalized signal for each 200 bp bin across the genome. Normalized signal is defined as the following:

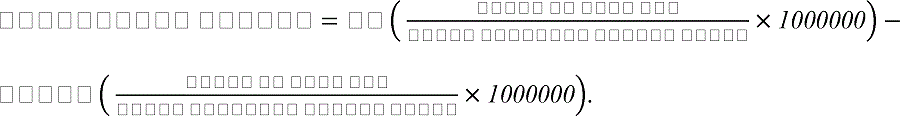

To assess the reproducibility of ChIP-Seq data, for a given histone modification mark, we divided the mouse genome into consecutive 1 kb windows. Normalized ChIP-Seq signals in windows were used for computing genome-wide correlation using Pearson correlation coefficient. Windows with zero signals in both replicates were removed before calculating correlation.

### RNA sequencing

Cells were sorted into TRIzol-LS (Invitrogen) and total RNA was purified using RNeasy micro kit (Qiagen). Total RNA was quantified with RNA HS Kit for Qubit fluorometer (Life Technologies) and analyzed for integrity using RNA 6000 Pico Kit for 2100 Bioanalyzer (Agilent). mRNA was purified from total RNA using NEBNext Poly(A) mRNA magnetic isolation module (NEB). Poly(A)-selected mRNA was fragmented to an average size of 300 nt, reverse transcribed to generate double-stranded cDNA, and converted to a paired-end sequencing library using NEBNext Ultra RNA Library Prep Kit for Illumina (NEB) according to the vendor’s instructions including the optional double size selection procedure using Agencourt AMPure XP beads (Beckman Coulter). Prepared libraries were quantified with dsDNA HS Kit (Life Technologies) for Qubit and the size distribution was assessed using High Sensitivity DNA Kit for Bioanalyzer. Libraries were sequenced on Illumina HiSeq 2500 in paired-end mode with the read length of 75 nt.

### RNA-Seq data processing and differential expression analysis

Sequencing reads were aligned to the mouse genome (mm9) using Tophat (Kim et al., 2013). Only uniquely mapped reads with fewer than 2 mismatches were used for downstream analyses. The Ensembl database (release 66) was used as the source of gene annotation. The featureCounts function of the Subread package (Liao et al., 2014) was used to extract both gene-level and transcript-level read counts. Differential expression analysis was performed using the edgeR algorithm (Robinson and Oshlack, 2010). ANOVA was first used to identify genes that are differentially expressed across any of the five populations with a False Discovery Rate (FDR) < 0.01. Next, pair-wise comparisons were conducted to identify genes differentially expressed between two populations with FDR < 0.05 and fold change > 2.

### Clustering of expression profile using STEM

Gene expression values were expressed as Fragment Per Kilobase of transcript per Million mapped reads (FPKM). Short-time series expression miner (STEM) (Ernst et al., 2005) was used for clustering with default parameter setting except that minimum correlation of profile was set to 0.9 to group similar expression profiles.

### Enhancer prediction using histone modification ChIP-Seq data

Enhancers were predicted using the CSI-ANN algorithm (Firpi et al., 2010). The inputs to the algorithm are normalized ChIP-Seq signals of four histone marks (H3K4me1, H3K4me3, H3K27ac, and H3K27me3). The algorithm combines signals of all histone marks and uses an artificial neural network-based classifier to make predictions.

### Identification of enriched TF binding motifs at enhancers

TF DNA binding motifs were downloaded from the Cis-BP database (Weirauch et al., 2014). Similar motifs were merged using TomTom (Bailey et al., 2009). Find Individual Motif Occurrences (FIMO) (Grant et al., 2011) was used to scan enhancer sequences defined by CSI-ANN. A *P*-value cutoff of 1E-5 (multiple testing corrected using Bonferroni’s method) was used to identify binding sites according to the TF motif model. For background, we used 9,500 enhancers from 19 non-hematopoietic cell lines reported by the ENCODE project (Shen et al., 2012) (brown adipose tissue, brain (E14.5), cerebellum, cortex, heart (E14.5), heart, intestine, kidney, limb (E14.5), liver (E14.5), liver, lung, mouse embryonic fibroblasts, mouse ESC, olfactory, placenta, spleen, testes, and thymus, Table S15). Hypergeometric distribution was used to compute the enrichment *P*-values. Benjamini-Hochberg method was used to correct *P*-values for multiple testing.

### Construction of transcriptional regulatory networks

We developed a computational pipeline to infer target genes of a TF based on the probabilities of enhancer, TF binding site, and enhancer-promoter linkage. The probability of an enhancer based on its histone mark signals, *P_enh_*, is defined by the CSI-ANN algorithm. The probability of an enhancer-promoter linkage, *P_EP_*, is defined by the IM-PET algorithm. To determine if a TF occupies an enhancer, we first use a compendium of 1147 mouse TF motifs curated in the Cis-BP database (Weirauch et al., 2014). We calculate the probability that a TF is bound to its site in the enhancer, *P_TFBS_*. Given an enhancer sequence *l* and a PSSM *M* representing a TF binding motif, the binding probability can be approximated as following according to (Stormo and Fields, 1998):

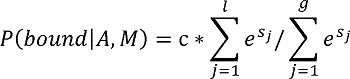

where *l* is the length of *A* and *g* is the length of the background sequence, *S_j_* is the score of the sequence word starting at position *j* according to the PSSM, and *c* represents the concentration of the TF in the cell. In our analysis, we will use the entire sequence of chromosome one as the background, and the mRNA expression level of the TF as an estimate of *c*. Finally, the probability of a TF regulating a target gene, *P_TG_*, is defined to be the product of the three probabilities, *P_TG_* = *P_enh_* x *P_TFBS_* x *P_EP_*.

### Identification of key transcription factors based on their regulatory potential

We used the constructed TRNs to identify key transcriptional factors for the EHT. To this end, we assume that key TFs are closer to the set of differentially expressed genes in the TRN, either via direct or indirect connections. Based on this assumption, we computed a distance between two genes, *i* and *j*, in the TRN as following:

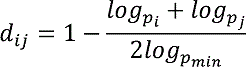

where *p_i_* and *p_j_* are the differential expression *P*-values for genes *i* and *j*, respectively. *p_min_* is the minimum differential expression *P*-value among all genes in the TRN. With the distance-weighted TRNs, we calculated an average shortest distance between a given TF and all differentially expressed genes in the network. Statistical significance of average shortest distance was computed using a null distribution based on the given TF and randomly selected genes. We selected the sets of key TF as follows. First we ranked the TFs based on their distance to differentially expressed genes in the TRN. Then, starting from the highest ranked TF, we iteratively computed the cumulative union of the regulons of TF sets, having *n* number of TFs, at *n*^th^ step. Next, we computed the difference between the expected regulon size with the actual regulon size. We set the cutoff for the number of key TFs at the inflection point of the curve.

### Overlap between public TF ChIP-Seq data and SP3 binding regions

We compared the predicted SP3 bound enhancers and promoters in our data and TF peaks from public ChIP-Seq data as follows. For the GSE40235 dataset (Lichtinger et al., 2012), we called the ChIP-seq peaks using MACS 1.4.2 with the parameter settings provided by the authors (mfold=10,16-bw=100-pvalue=1e-9-gsize=1,870,000,000). For the GSE69099 dataset (Goode et al., 2016), we used the TF peaks provided by the authors. We first identified the target genes of SP3 in each TRN, then extracted the promoters and enhancers that target those genes and had a significant binding site hit for SP3 (FIMO *P*-value < 1e-5). We then identified ChIP-Seq peaks overlapping with SP3 bound enhancers and promoters in our dataset. We used the hypergeometric test to assess the statistical significance of the overlap using the total number of regulatory regions (enhancers and promoters) for each developmental transition and the total number of peaks for each TF as the background.

### Identification of mouse orthologs in zebrafish

Orthologs of *Sp3* and *Maz* in zebrafish were identified using four databases: MGI (http://www.informatics.jax.org), ZFIN (https://zfin.org), OrthoRetriever (http://lighthouse.ucsf.edu/orthoretriever) and DIOPT (http://www.flyrnai.org/diopt).

### CRISPR-Cas9 mediated gene knockout in zebrafish

Genes were knocked out with Alt-R™ CRISPR-Cas9 crRNA from Integrated DNA Technologies (IDT) following the manufacturer’s instructions. Briefly, crRNAs were designed using CRISPOR (http://crispor.tefor.net/). To generate mutants, 900 picoliter of 7.5 µM ribonucleoprotein complex (7.5 µM crRNA and 7.5 µM Cas9 enzyme (IDT)) was injected directly into 1-cell-stage embryos (strain Casper) and up to 200 zebrafish embryos per target gene. Editing efficiency was measured using the TIDE assay. The mRNA and protein levels of target genes were measured using RT-qPCR and Western Blot, respectively. Primer sequences are included in Table S14.

### Tracking of indels by decomposition (TIDE)

Genomic DNA was extracted from embryos using the HotSHOT method(Meeker et al., 2007). A short stretch of genomic region (∼600-800 bp) flanking the target site was amplified using Q5 Hot Start High Fidelity 2 × Master Mix (NEB). Amplified sequences were purified with the QIAquick PCR purification kit (Qiagen) and sequenced by Sanger sequencing. A fragment amplified using wild type zebrafish was used as a control. Primer sequences are included in Table S14.

### Whole-mount in situ hybridization (WISH) in zebrafish

Zebrafish embryos were dechorionated using pronase, fixed in 4% paraformaldehyde at 4 °C overnight, dehydrated in methanol and stored at −20 °C. Both sense and anti-sense RNA probes were synthesized from restriction enzyme linearized plasmids transcribed with T7 or SP6 polymerase in the presence of RNA DIG labelling mix (Roche). Newly synthesized probes were purified using mini QuickSpin columns according to the manufacturer’s protocol (Roche). Synthesis of RNA probes was performed as described (North et al., 2007; Song et al., 2004). WISH was performed using an established protocol (Thisse and Thisse, 2008) with some modifications. Briefly, embryos stored in methanol were rehydrated in successive dilutions of methanol in 1× PBS. After washing in 1× PBST, embryos were permeabilized by digestion with 10 µg/mL proteinase K at room temperature. Digestion with proteinase K was stopped by 4% paraformaldehyde in 1× PBS and embryos were washed in 1× PBST. After prehybridization at 65 °C, the embryos were hybridized in a 70 °C hybridization oven overnight. Following several washes, embryos were incubated with anti-DIG antibody at 4 °C overnight. Embryos were then washed, stained and mounted in 100% glycerol for imaging using a Leica S8 APO stereo microscope with a Leica DFC420 camera. At least 20 embryos per condition were scored.

### Whole-mount immunofluorescence and confocal microscopy

Embryos were prepared as described previously (Yokomizo et al., 2012). The following primary antibodies were used: rat anti-mouse CD117 (eBioscience, 14-1171-85, 1:250), rat anti-mouse CD31 (BD Pharmingen, 557355, 1:500) and rabbit anti-human/mouse Runx (Abcam, ab92336, 1:250). Secondary antibodies were goat anti-rat Alexa Fluor 647 (Molecular Probes, A-21247, 1:500), goat-anti rat Alexa Fluor 555 (Molecular Probes, A-21434, 1:1000) and goat anti-rabbit Alexa Fluor 488 (Molecular Probes, A-11034, 1:1000). Images were acquired on a Zeiss LSM 710 AxioObserver inverted microscope with ZEN 2011 software. The Zeiss LSM 710 is equipped with 488, 543 and 633 nm wavelengths. Images were processed with Fiji software (Schindelin et al., 2012).

### Analysis of blood vessel diameter in zebrafish

Blood vessel diameter analysis was done using 48 hpf zebrafish embryos. For dorsal aorta (DA) and posterior cardinal vein (PCV), measurements were taken at the midway point between intersegmental vessels (ISV) along the yolk extension, and the mean was used as an average diameter per embryo. For ISV, 4 measurements were made along each ISV between the DA and the dorsal longitudinal anastomotic vessel (DLAV), and the mean was used as an average diameter per ISV.

